# ccnWheat: A Database for Comparing Co-expression Networks Analysis of Allohexaploid Wheat and Its Progenitors

**DOI:** 10.1101/2022.01.17.476536

**Authors:** Zhongqiu Li, Yiheng Hu, Xuelian Ma, Lingling Da, Jiajie She, Yue Liu, Xin Yi, Yaxin Cao, Wenying Xu, Yuannian Jiao, Zhen Su

**Author notes:** Corresponding authors., (Su Z), (Jiao Y).

## Abstract

Genetic and epigenetic changes after polyploidization events could result in variable gene expression and modified regulatory networks. Here, using large-scale transcriptome data, we constructed co-expression networks for diploid, tetraploid, and hexaploid wheat species, and built a platform for comparing co-expression networks of allohexaploid wheat and its progenitors, named ccnWheat. ccnWheat is a platform for searching and comparing specific functional co-expression networks, as well as identifying the related functions of the genes clustered therein. Functional annotations like pathway, gene family, protein-protein interactions, microRNA (miRNA), and several lines of epigenome data are integrated in this platform, and Gene Ontology (GO) annotation, gene set enrichment analysis (GSEA), motif identification, and other useful tools are also included. Using ccnWheat, we found that the network of *WHEAT ABERRANT PANICLE ORGANIZATION 1 (WAPO1)* has more co-expressed genes related to spike development in hexaploid wheat than its progenitors. We also found a novel motif of CArG specifically in the promoter region of *WAPO-A1*, suggesting that neofunctionalization of the *WAPO-A1* gene affects spikelet development in hexaploid wheat. ccnWheat is useful for investigating co-expression networks and conducting other associated analyses, and thus facilitates comparative and functional genomic studies in wheat. ccnWheat is freely available at http://bioinformatics.cau.edu.cn/ccnWheat.

## Introduction

Wheat, in the Poaceae family, is the most widely grown food crop worldwide, providing an important source of nutrients for millions of people. Global food demand is increasing rapidly, with a 60%-70% increase in food production required by 2050 [1]. Basic research and optimization of wheat breeding is necessary to meet this demand. However, the bread wheat genome is large and complex (16 GB) [2], and thus wheat research lags behind that on rice and maize. This lag is mainly because bread wheat is a recent allohexaploid, formed via two consecutive allopolyploidization events. The diploid wheat *Triticum urartu* (AA) and a yet unknown *Aegilops* species formed tetraploid wheat *Triticum dicoccoides* (AABB) around 0.5 million years ago (hereafter MYA). Subsequently, tetraploid wheat and *Aegilops tauschii* (DD) hybridized to form the hexaploid wheat *Triticum aestivum* (AABBDD) around 0.01 MYA [3, 4]. As wheat is a primary food crop, wheat scientists have worked to sequence the genome of hexaploid wheat and its progenitors, and the completion of genome sequencing has laid a solid foundation for studying functional, comparative, and evolutionary genomics of wheat [5, 6].

Currently, public databases for wheat can be divided into four types: Genome, Transcriptome, Proteome, and other. Genome databases, like the Wheat@URGI portal [7], GrainGenes [8], CerealsDB [9], Wheat-SnpHub-Portal [10], WheatGmap [11], and Triticeae-GeneTribe [12] provide genomic or genetic data and other useful tools. Transcriptome databases including expVIP [13] and Wheat eFP Browser (http://bar.utoronto.ca/efp_wheat/cgi-bin/efpWeb.cgi), are usually used to provide the expression patterns of homeologs. Transcriptome-based co-expression networks like WheatNet have been constructed by DNA microarray datasets with an early genome assembly version [14]. Knetminer [15] and WheatOmics [16] only contain a hexaploid wheat network, not including wheat progenitors. Wheat Proteome provides searchable organ and developmental stage proteomic data [17]. Other databases like the Triticeae Toolbox (T3) has phenotype and genotype data for barley, wheat, and oat [18]. Wheat microRNA Portal has integrated the abiotic stress response miRNA in wheat [19]. Despite the accumulation of large-scale RNA-Seq data and improved and high-quality wheat genomic sequence, there are still gaps in meeting demands for co-expression analysis within allohexaploid wheat and its progenitors.

Thus, we developed the co-expression network comparison database ccnWheat for allohexaploid wheat and its progenitors, including four global networks (*T. aestivum, T. dicoccoides, T. urartu*, and *Ae. tauschii*) and two *T. aestivum* conditional networks (tissue-specific and stress-treated). ccnWheat integrates functional and epigenome sequencing data, and includes useful tools like gene set enrichment analysis (GSEA), Gene Ontology (GO) analysis, and motif analysis, which will help bench scientists easily pick key candidate genes for functional studies. In addition, this database will provide genomic scientists a useful source for deciphering key molecular modules during the formation, evolution, and domestication of wheat.

## Construction and content

### Data sources and processing

For genome data, *T. aestivum* (AABBDD) was based on the International Wheat Genome Sequencing Consortium (IWGSC) Chinese Spring v1.0 genome assembly and v1.1 annotation (URGI) [2]; *T. dicoccoides* (AABB) was based on the Zavitan WEW_v1.0 genome assembly and annotation [20]; *T. urartu* (AA) was based on the IGDB (Institute of Genetics and Developmental Biology, Chinese Academy of Sciences) genome assembly and annotation [21]; and *Ae. tauschii* (DD) was based on the CAAS (Chinese Academy of Agricultural Sciences) genome assembly and annotation [22].

For transcriptome data, we used most of the available RNA-seq datasets in addition to the common tissues of leaf, root, and grain in the four studied species for robust constructed co-expression networks [23, 24]. Finally, we collected 425 transcriptomic datasets (112 for *T. aestivum*, 153 for *T. dicoccoides*, 90 for *T. urartu*, and 70 for *Ae. tauschii*) from the NCBI Sequence Read Archive (SRA) (for detailed sample information see Table S1). Quality control was conducted based on FastQC software (http://www.bioinformatics.babraham.ac.uk/projects/fastqc/) and Trimmomatic software [25] was used to remove adapter sequences and low-sequencing-quality bases. The remaining sequence data (112 for *T. aestivum*, 153 for *T. dicoccoides*, 76 for *T. urartu*, and 42 for *Ae. tauschii*) were mapped to corresponding genomes, then the fragments per kilobase of transcript per million mapped reads (FPKM) values of all protein-coding genes were calculated from each sample with parameter settings --star and --estimate-rspd using RSEM [26]. Those samples with mapping rate < 50% were filtered out. Then, the R package “pheatmap” (https://github.com/raivokolde/pheatmap) was used to perform cluster analysis on all datasets, and the outlier samples were excluded (Figure S1 and S2). According to the boxplot of the reading score distribution, the quality of the remaining samples was acceptable (Figure S3). The mapping results of these data are listed in Table S2.

For epigenomic data, all 25 *T. aestivum* epigenomic datasets including H3K4me3, H3K4me1, H3K36me3, H3K27me3, H3K9me2, H3K9ac, H3K27ac, DNase-seq, and CENH3 were downloaded from public platforms, including the NCBI Sequence Read Archive (SRA). Quality control was conducted based on trim_galore (https://www.bioinformatics.babraham.ac.uk/projects/trim_galore/) software with parameter settings q = 25, stringency = 3. The sequence reads were mapped to the *T. aestivum* accession Chinese Spring v1.0 reference genome with the maximal exact matches (MEM) algorithm and default parameters by BWA software [27]. The enriched regions were called by MACS software [28] with the nomodel parameter. The details and mapping results of these data are listed in Table S3.

For collinear ortholog pairs, we first computed the top 5 BLAST results between two species using protein sequences based on the rank of bit score and that met an E-value threshold of 10^-5^, which were suggested by MCScanX tools [29]. Then the top 5 BLAST results with gff (general feature format) files were used to establish collinear ortholog pairs in *T. urartu, Ae. tauschii, T. dicoccoides*, and *T. aestivum* through MCScanX tools with default parameters.

### Co-expression network construction and network comparison

We used the calculated FPKM values for constructing the four global networks and two *T. aestivum* conditional (tissue-specific and stress-treated) networks based on the Pearson correlation coefficient (PCC) and mutual rank (MR) algorithm [30]; every network covered at least 84.70% of genes (**Table 1**). We used the PCC and MR to measure the co-expression relationship between genes. First, based on the PCC distribution diagram of all gene pairs, the thresholds for negative and positive correlation were the values of the lowest 5% PCC (−0.25) and highest 5% PCC (0.5) in the *T. aestivum* global network, respectively (Figure S4). Second, we used the MR method to exclude poor co-expression gene pairs, because MR has been used successfully for this purpose in several plants such as *Arabidopsis*, maize, and bamboo [31]. Furthermore, the GO terms of biological process related to multiple genes in the interval [4, 20] were used to evaluate the accuracy of the co-expression network through the receiver operating characteristic (ROC) curve [32]. Finally, to compare the *T. aestivum* global networks and conditional networks or other species’ networks while considering each network’s coverage and connectivity, we set the MR of all networks to 30.

**Table 1.** Information about the four wheat species network.

| Co-expression network | AABBDD global | AABBDD tissue-specific | AABBDD stress-treated | AABB | AA | DD |
| --- | --- | --- | --- | --- | --- | --- |
| Nodes | 103,512 | 102,959 | 94,849 | 56,858 | 36,613 | 36,286 |
| (nodes/all genes %) | (95.9%) | (95.4%) | (87.9%) | (87.5%) | (88.2%) | (84.70%) |
| Positive edges | 985,277 | 1,004,338 | 892,249 | 564,671 | 366,625 | 392,654 |
| Negative edges | 418,096 | 459,775 | 388,565 | 274,275 | 160,254 | 164,741 |
| Sample count<br>(excluding duplicates samples) | 112<br>(49) | 61<br>(32) | 51<br>(17) | 153<br>(20) | 76<br>(41) | 42<br>(20) |
| Tissues | Spike,<br>rachis,<br>ovary,<br>**root,**<br>**grain,**<br>**leaf,**<br>crown,<br>seedling,<br>seed, stem,<br>flag leaf | Spike,<br>rachis,<br>ovary,<br>**root,**<br>**grain,**<br>**leaf,**<br>crown,<br>seedling,<br>seed, stem,<br>flag leaf | **Leaf,**<br>**grain,**<br>**root,**<br>crown | Spike,<br>**leaf,**<br>**root,**<br>**grain,**<br>flower,<br>glume,<br>lemma | Shoots,<br>**root, leaf,**<br>**endosperm** | Spikes,<br>seedling,<br>**grain,**<br>**endosperm,**<br>**leaf,**<br>**root,**<br>shoots,<br>immature<br>seeds |
| Treatment | Heat,<br>drought,<br>chitin,<br>Flg22 | - | Heat,<br>drought,<br>chitin,<br>Flg22 | - | Bgt, Cd,<br>cold | - |
| AUC | 0.8599 | 0.8386 | 0.8081 | 0.7486 | 0.711 | 0.579 |
| (cutoff) | (MR30) | (MR30) | (MR30) | (MR30) | (MR30) | (MR30) |
| Module (number) | K = 9 | K = 7 | K = 10 | K = 10 | K = 10 | K = 8 |
|  | (1867) | (3913) | (1118) | (625) | (524) | (851) |

*Note:* AUC, Area Under Curve; The tissues marked in bold represent the RNA-seq data of these tissues were found in all four studied species.

Further, in order to verify whether the networks we built are comparable among allohexaploid wheat and its progenitors, we collected genes from the literature (such as light-dependent chlorophyll accumulation gene *TaCHLH* [33], peroxidase gene *TaPOD1* [34], *Storage protein activator (TaSPA)* or Photosynthesis pathway (*LHCA2, LHCA3, LHCB7*, and *psaK*)) (Table S4). We queried the expression values of these genes in the leaf, root, and grain of the four species. The results suggested that the expression trends of these genes in samples with different developmental stages or stress treatments are still consistent and comparative among four species for analyzing the network (Figure S5).

### Functional modules identification

We used the CFinder software to identify modules containing more densely connected genes by combining positive and negative gene pairs together [35] (Table 1). Then we used the gene set annotations, like gene families, GO, and metabolic pathways (detailed information is in the Functional annotation and gene annotation section) to predict the functions of modules. Non-significant entries were then filtered out using Fisher’s tests and the multiple test correction method “Benjamini-Yekutieli” (false discovery rate (FDR), as referred to in the PlantGSEA [36]). As a result, 1867 functional modules in AABBDD, 625 functional modules in AABB, 524 functional modules in AA, and 851 functional modules in DD were obtained. These modules may be related to important agronomic traits.

### Usage of co-expression network tools

In ccnWheat, a network search tool for one gene or a gene list was provided for the four global and two conditional (tissue-specific and stress-treated) co-expression networks, which were visualized using Cytoscape [37] (**Figure 1A**). Furthermore, we established collinear ortholog pairs in AA, DD, AABB, and AABBDD using MCScanX [29] tools. We used collinear gene pairs to determine the correlations between diploid and polyploid wheat. With the relationship of homologous genes, three types of co-expression networks between polyploid wheat and its progenitors can be compared: genes of a single species can be compared with global network and tissue specific network or stress treatment network in the network comparison tool (Figure 1B); genes of two species, such as AA vs AABB, DD vs AABBDD, AABB vs AABBDD in the network comparison tool; and genes of three species (DD, AABB, AABBDD) in the ortholog network comparison tool. For instance, users can submit one gene of interest in our ortholog network comparison tool, and choose two or three species for the orthologous pair of this gene. Then all the orthologous gene pairs are highlighted and linked to each other with brown lines in the network to exhibit the conservation and diversification of the regulatory network during wheat evolution (Figure 1C). For all genes in the network, the annotations and relationships of genes in a network are listed in tables. GSEA, GO analysis, and motif analysis tools are used to find the potential function and regulation in the promoter region of genes in the network (Figure 1D) as well as the expression profile and the distribution of genes on chromosomes of the network (Figure 1E). Moreover, the gene expression profiles and cis-elements of the homeologous sub-networks can be compared to find the similarities and differences of the network.

**Figure 1.**
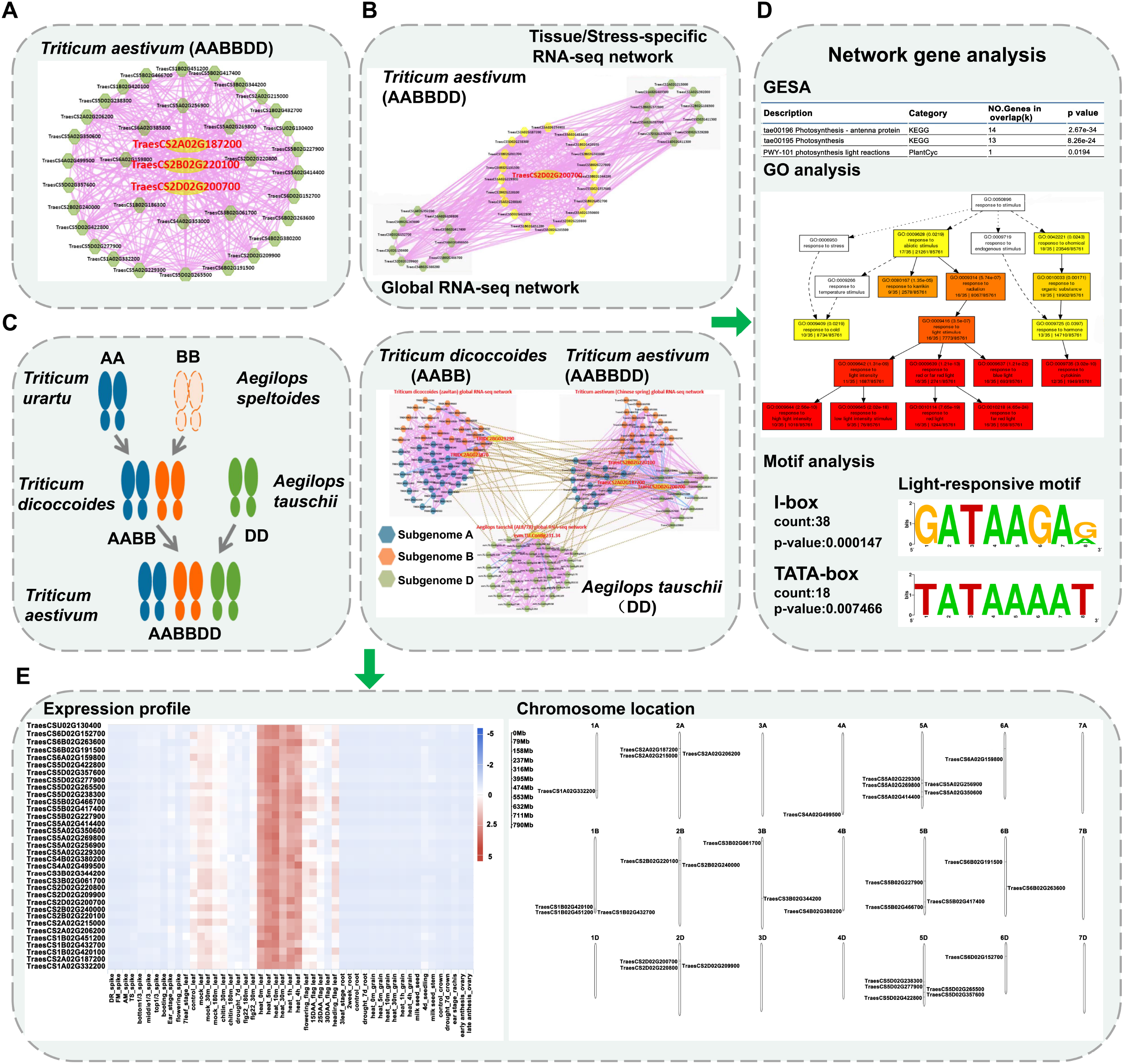
Description of networks in the database. **A.** An example of gene search results in *T. aestivum* (AABBDD). The largest red node represents the gene queried; the green node represents the co-expressed gene of the queried gene. The pink edge links two genes that have a positive co-expression relationship; the blue edge links two genes that have a negative co-expression relationship. **B.** Global RNA-seq network vs tissue-specific RNA-seq network in *T. aestivum* (AABBDD). The biggest red node represents the gene queried; the yellow node represents the overlapping co-expressed genes between two networks; the green node in the gray box represents specific genes in their respective networks. **C.** Network comparison of *T. aestivum* (AABBDD), *T. dicoccoides* (AABB), *T. urartu* (AA), and *Ae. tauschii* (DD). A brown colored edge links two genes with an orthologous relationship between two species; blue nodes represent the co-expressed genes of subgenome A; orange nodes represent the co-expressed genes of subgenome B; green nodes represent the co-expressed genes of subgenome D. **D.** Results of analysis tools that we provided. GSEA, GO, and motif analysis can be performed directly on the network results page. **E.** Expression profiling of all genes in the network displayed by heatmap. The chromosome positions of all genes are also displayed. DR, double-ridge stage; FM, floret meristems; AM, anther primordia stage; TS, tetrads stage; DAA, days after anthesis.

### Functional annotation and gene annotation

For functional annotation (**Figure 2**A and **Table 2**), there are five types of data in ccnWheat, which can be searched by users to predict gene function. On the one hand, integrating these data is of great significance for genome-level gene annotation; on the other hand, these functional annotations can be used as background gene sets for data mining, such as enrichment analysis for network or functional modules.

**Figure 2.**
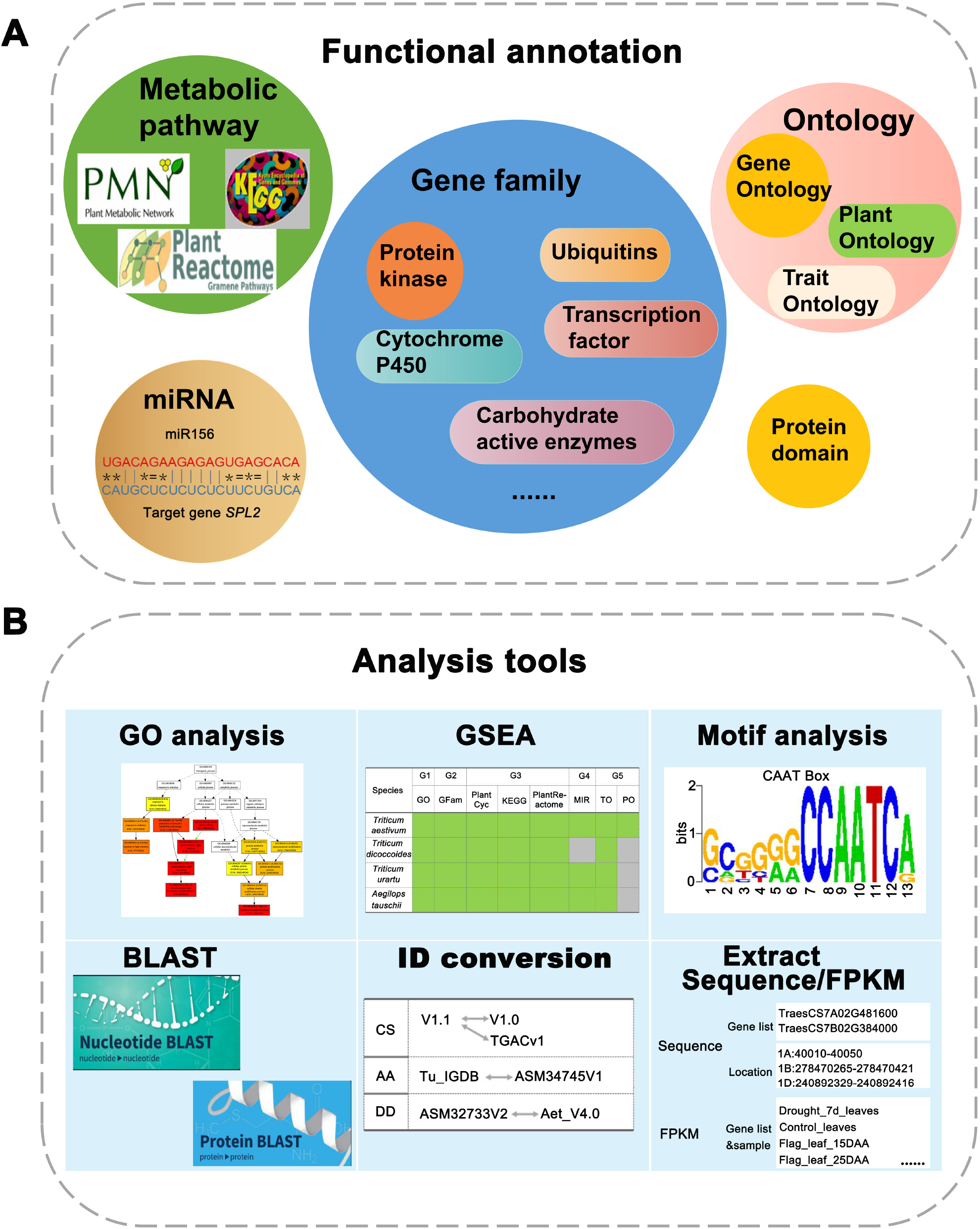
Description of functional annotation and support tools in the database. **A.** Five types of data are included in Functional annotation. Gene families include ubiquitin, transcription factor/regulator, protein kinase, cytochrome P450, carbohydrate-active enzymes, and so on. Ontology (such as GO, PO, TO), Protein domain, MicroRNA, Metabolic pathways including PlantCyc, KEGG, and plant reactome, can be browsed. **B.** Tools, like GSEA analysis, GO analysis, motif analysis, BLAST, ID conversion, Extract sequence and FPKM, UCSC genome browser (including epigenome data in *T. aestivum* (AABBDD), and RNA-seq sample in *T. dicoccoides* (AABB) (not shown in the picture) are supported in ccnWheat.

**Table 2.** Functional annotation collected from public databases or annotated by public software.

| Classification | Category<br>(source/method) | AABBDD | AABB | AA | DD |
| --- | --- | --- | --- | --- | --- |
|  |  | module | module | module | module |
|  |  | count | count | count | count |
|  |  | (gene) | (gene) | (gene) | (gene) |
| Gene | GO | 7391 | 9716 | 1814 | 7612 |
| Ontology | (IWGSC/Blast2GO/MBKBASE) | (85,762) | (49,242) | (21,216) | (24,143) |
| Trait | TO | 276 (1000) | 254 | 232 | 262 |
| Ontology | (IWGSC/ortholog relationship) |  | (528) | (218) | (308) |
| Plant | PO | 444 | - | - | - |
| Ontology | (IWGSC) | (63,687) |  |  |  |
| Pfam domain | Function domain | 4021 | 4052 | 3675 | 3930 |
|  |  | (83,690) | (49,301) | (26,636) | (26,275) |
| Gene family | Ubiquitin | 20 | 20 | 20 | 19 |
|  | (UUCD) | (6641) | (2667) | (1503) | (2068) |
|  | Transcription factor/regulator | 90 | 63 | 90 | 91 |
|  | (iTAK) | (7913) | (4622) | (2325) | (2337) |
|  | Protein kinase | 125 (5217) | 126 | 120 | 122 |
|  | (iTAK) |  | (3126) | (1623) | (1183) |
|  | Cytochrome P450 |  | 242 | 211 | 232 |
|  | (BLAST+Pfam) | 264 (1564) | (823) | (462) | (489) |
|  | carbohydrate-active enzyme | 94 | 93 | 91 | 92 |
|  | (ortholog) | (4290) | (2749) | (1326) | (1456) |
| Metabolic | Plant Reactome |  | 204 | 192 | 202 |
| pathway | (gramene) | 204 (3478) | (2063) | (654) | (976) |
|  | PlantCyc |  | 494 | 426 | 434 |
|  | (PlantCyc/ortholog) | 494 (7100) | (4209) | (2057) | (2228) |
|  | KEGG | 139 | 139 | 112 | 140 |
|  | (KEGG/KAAS/MBKBASE) | (15,528) | (9312) | (2246) | (4692) |
| miRNA | miRNA target |  |  |  |  |
|  | (miRbase/IWGSC/ literature) | 9626 | - | 1702 | 173 |
| PPI | protein-protein interaction | 149,844 | 53,169 | 16,726 | 16,226 |
|  | (Inparanoid) | (12,972) | (9452) | (4818) | (5105) |

Ontology has become a critical tool for enabling cross-species comparative analyses and increasing data sharing and reusability [38]. So we collected 7391 GO, 444 Plant Ontology (PO), and 276 Plant Trait Ontology (TO) annotation entries of AABBDD from URGI (https://wheat-urgi.versailles.inra.fr/Data); the 1814 GO annotation entries of AA were from MBKBASE (http://www.mbkbase.org/Tu/); and the 7612 and 9716 GO respective annotation entries of DD and AABB were annotated by Blast2GO software [39].

Metabolic pathways are responsible for the biosynthesis of complex metabolites, having an impact on the growth and development of plants or aiding plants to respond to biotic and abiotic stress [40]. We collected Plant Reactome pathways of all species from Gramene [41]; PlantCyc pathways of AA, DD, and AABBDD were integrated from PlantCyc [42], while AABB was predicted by orthologs. For the KEGG pathways (https://www.kegg.jp/kegg/), DD came from KEGG using the ID conversion; AABBDD and AABB KEGG pathways were annotated by the KEGG tool KAAS; and AA came from MBKBASE (http://www.mbkbase.org/Tu/).

Protein domains are an important part of proteins, and many domains either have specific functions or contribute to the function of their proteins in a specific way [43]. Protein domains were annotated by PfamScan tools (https://www.ebi.ac.uk/Tools/pfa/pfamscan/) based on a hidden Markov model [44]. We ultimately identified 4021 functional domains with 83,690 genes in AABBDD; 4052 functional domains with 49,301 genes in AABB; 3675 functional domains with 26,636 genes in AA; and 3930 functional domains with 26,275 gene in DD. Users can submit a gene list to ccnWheat, and then the protein domains and detailed information can be extracted to proceed to the next analysis (like adding protein domains when constructing evolutionary trees).

Gene families such as transcription factors (TFs)/transcriptional regulators (TRs), protein kinases (PKs), carbohydrate-active enzymes, and ubiquitin, play important biological roles. We used iTAK software (http://bioinfo.bti.cornell.edu/cgi-bin/itak/index.cgi) to identify TFs and PKs. We obtained a hidden Markov model from UUCD (http://uucd.biocuckoo.org/) to identify ubiquitin families. InParanoid (http://inparanoid.sbc.su.se/cgi-bin/index.cgi) software and Pfam domains were used to predict the carbohydrate-active enzyme families. So we annotated 20 ubiquitin families with 6,641 genes, 90 TF/TR families with 7913 genes, 125 protein kinase families with 5217 genes, and 94 carbohydrate-active enzyme families with 4290 genes in AABBDD; 20 ubiquitin families with 2667 genes, 63 TF/TR families with 4622 genes, 126 protein kinase families with 3126 genes, and 93 carbohydrate-active enzyme families with 2749 genes in AABB; 20 ubiquitin families with 1503 genes, 90 TF/TR families with 2325 genes, 120 protein kinase families with 1623 genes, and 91 carbohydrate-active enzyme families with 1326 genes in AA; and 19 ubiquitin families with 2068 genes, 91 TF/TR families with 2337 genes, 122 protein kinase families with 1183 genes, and 92 carbohydrate-active enzyme families with 1456 genes in DD. By comparing the protein sequences of hexaploid AABBDD and its progenitors with wheat protein sequences of CYP450s (http://drnelson.uthsc.edu/cytochromeP450.html), we further identified their P450 domains using PfamScan (https://www.ebi.ac.uk/Tools/pfa/pfamscan/). Finally 1564, 823, 462, and 489 full-length CYP450 genes were found in AABBDD, AABB, AA, and DD, respectively (For detailed information and related analysis about CYP450s see File S1).

MiRNA functions in plant developmental plasticity, abiotic/biotic responses, and symbiotic/parasitic interactions [45]. Integrated miRNA data including mature miRNA and precursor miRNA, 9626 AABBDD’ miRNA and 173 DD’ came from miRbase [46] and IWGSC; 1702 AA’ miRNA were collected from the literature [47], and all miRNA sequences were mapped to the corresponding genome by GMAP (http://research-pub.gene.com/gmap/), then the mature targets were predicted by psRNAtarget (http://plantgrn.noble.org/psRNATarget/). We also predicted protein-protein interaction (PPI) pairs in all species with some experimentally validated PPI pairs in *Arabidopsis* predicted by InParanoid [48]. There were 149,844 PPI with 12,972 genes in AABBDD; 53,169 PPI with 9452 genes in AABB; 16,726 PPI with 4818 genes in AA; and 16,226 PPI with 5105 genes in DD (Table 2).

For gene annotation, gene search results included all known and predicted information. Taking *WHEAT ABERRANT PANICLE ORGANIZATION 1 (WAPO-A1)* as an example, the gene detail page includes basic information: the gene locus is TraesCS7A02G481600; it is a 1457 bp gene with two exons located on chromosome 7A that encodes an F-box-like protein, related to spikelet number per spike; the best orthologous gene can be linked to the detailed function in *Arabidopsis thaliana* and *Oryza sativa*; and the gene, CDS, and protein can also be downloaded. In addition, the co-expression network is shown including orthologous genes and networks (global network, tissue and stress specific network) linked to the corresponding functional interface in diploid and polyploid wheat. Heuristic information, such as F-box ubiquitin family and GO terms related to flower development, regulation of circadian rhythm and so on, is displayed. Moreover, the details show an F-box-like functional domain with alignment start and end information; gene expression profiling in samples (*WAPO-A1* is specifically expressed in spike); and *WAPO-A1* included in the module shows the possible functional clues identified by CFinder. For histone modification, the University of California at Santa Cruz (UCSC) [49] genome browser shows obvious peaks for *WAPO-A1* in H3K4me3 and H3K27me3, which are related to spikelet or flower development (**Figure 3**).

**Figure 3.**
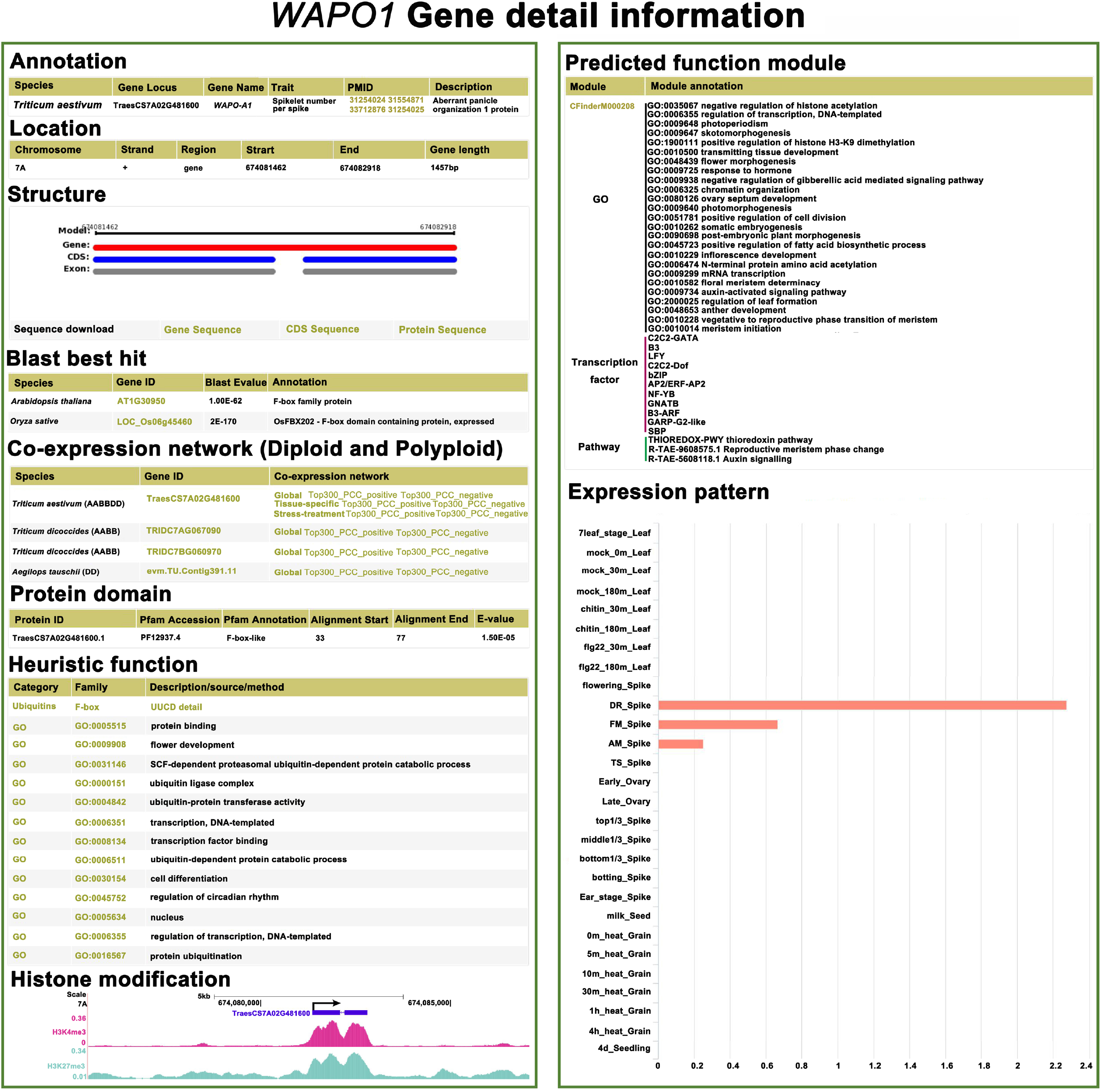
Details of gene annotation. Gene search results interface on the website, including gene annotation, gene location, structure, BLAST results, protein domain, co-expression network, heuristic function and predicted function module, expression pattern, and histone modification. The light yellow items link to the download page or detail page. DR, double-ridge stage; FM, floret meristems; AM, anther primordia stage; TS, tetrads stage.

### Supported analysis and tools

There are three category analysis tools, including GSEA, GO analysis, and cis-element enrichment. GSEA was based on the data analysis processing of PlantGSEA [36]. Here, we used functional data information as gene sets. The annotation entries with FDRs < 0.05 were used and displayed. GO analysis was based on agriGOv2 [50] data processing. Cis-element analysis (such as the functions of sequence scan, gene name scan, and custom scan) can identify significantly enriched motifs in the promoter regions of one gene and thus predict possible functions. We also provided BLAST (DNA and protein), ID conversion (in different genome versions), Sequence (gene ID or chromosome position), and FPKM extraction tools for users to conveniently obtain the information of a gene (Figure 2B).

### Case study: Function analysis of known gene *WAPO1* using ccnWheat

Polyploidy (*i.e.*, whole genomic duplication) is an important evolutionary feature in the plant kingdom, particularly in flowering plants [51], after which individual genes may experience nonfunctionalization, neofunctionalization, or subfunctionalization [52]. For example, *WAPO1* is an orthologue of rice gene *ABERRANT PANICLE ORGANIZATION 1* (*APO1*) and *Arabidopsis* gene *UNUSUAL FLORAL ORGANS* (*UFO*). *UFO* acts synergistically with floral meristem identity factor *LEAFY* (*LFY*) and restricts the expression of the class B floral organ identity genes in *Arabidopsis* [53]. The interactions between the orthologs of *LFY* and *UFO* have also been demonstrated in rice, petunia, *Antirrhinum majus*, and pea, suggesting that *LFY* and *UFO* are conserved among species [53, 54]. *EVERGREEN (EVG)* encodes a WOX homologous domain protein, which is only expressed in the initial lateral IM and participates in the activation of the *UFO* homologous gene *DOUBLETOP* (*DOT*) in *petunia*. The *EVG* ortholog of *COMPOUND INFLORESCENCE (S)* and *UFO* ortholog *ANANTHA (AN)* have similar effect on inflorescence meristems in tomato and related nightshades [53].

In wheat, *WAPO-A1* showed a highly significant association with total spikelet number (TSN) [55–58]. There are three genes (TraesCS7A02G481600 (*WAPO-A1*), TraesCS7B02G384000 (*WAPO-B1*), TraesCS7D02G468700 (*WAPO-D1*)) in AABBDD; two genes (TRIDC7AG067090 (not expressed), TRIDC7BG060970) in AABB; no gene in AA and one gene (evm.TU.Contig391.11) in DD. We searched the co-expression network of *WAPO1* using ccnWheat (**Figure 4**), and found that spike development related gene *wheat FLO/LFY WFL* [59], *TaSPL14* [60], *TaSPL17* [61], and *WUSCHEL-related homeobox TaWOX4/TaWUS* [62, 63], were directly co-expressed with *WAPO1* in AABBDD, but were not found in DD and AABB co-expression networks, even in the top 300 co-expressed genes. *WAPO-A1* also seems to be co-expressed with more genes related to spike development than *WAPO-B1* and *WAPO-D1. WAPO-A1b* in H2 haplotype (present in the Chinese Spring IWGSC v1.0 genome) was associated with higher spikelet number per spike (SNS) than *WAPO-A1c&d* in H3 haplotype (present in *WAPO-B1* and *WAPO-D1*) and *WAPO-A1a* in H2 haplotype [57]. *WAPO-B1* also have alleles conferring higher spikelet number at chromosome 7B [58]. So taken together, in addition to the effect of natural gene variation on gene expression, the co-expressed genes may also regulate spike development together with *WAPO1* in wheat.

**Figure 4.**
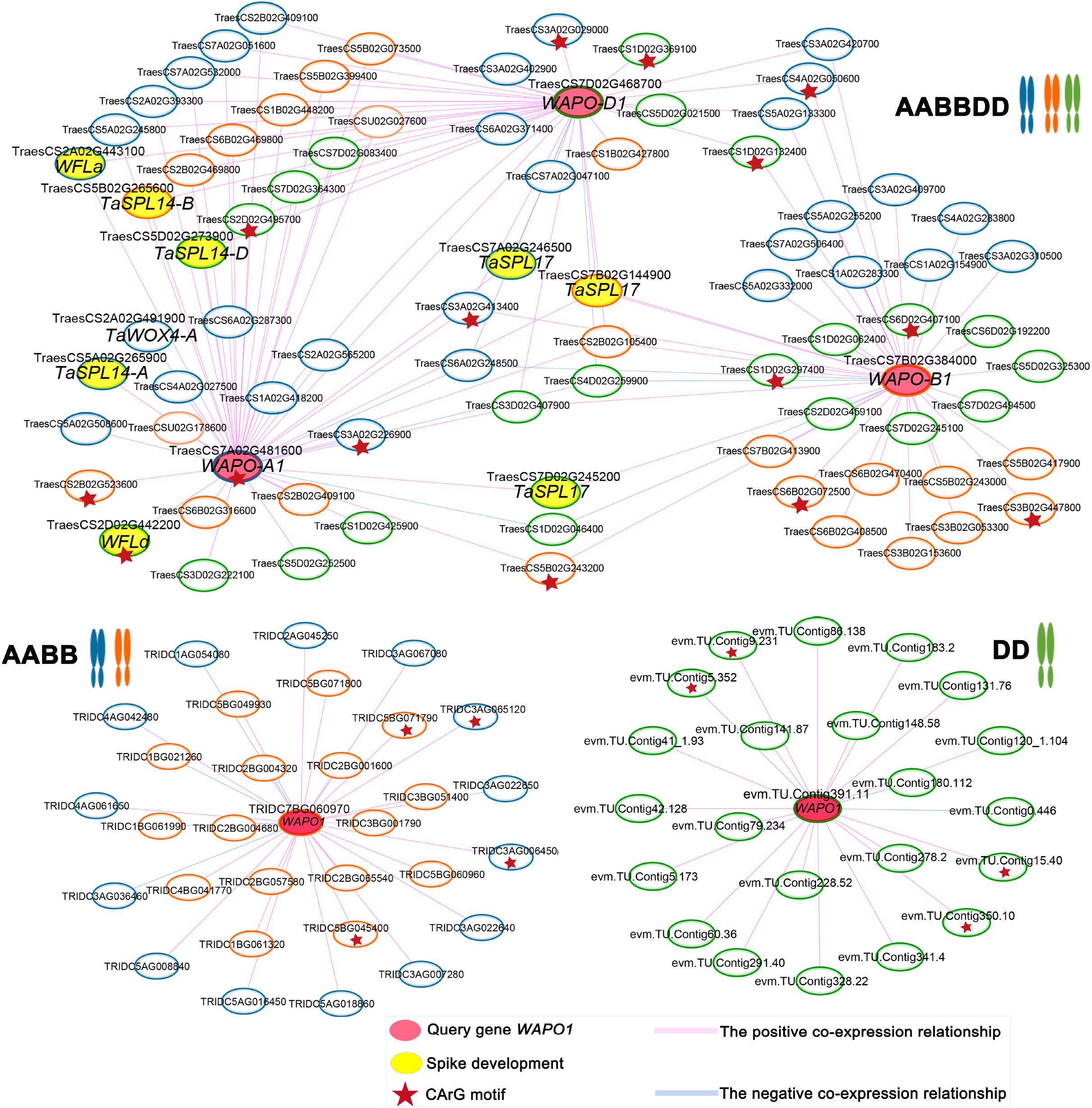
Global co-expression network analysis of *WAPO1*. Co-expression network of the *WAPO1* gene in DD, AABB, and AABBDD. The red node represents the query gene *WAPO1* and the yellow node represents spike development related gene. The blue circle represents the subgenome A gene; the orange circle represents the subgenome B gene; the green circle represents the subgenome D gene; and the red star means the CArG motif is included in the 3 kb upstream region of those genes.

GO analysis of the *WAPO1* co-expression network in AABBDD showed that these genes were related to reproductive shoot system development and post-embryonic development (Figure S6A). GSEA results for the network (Figure S6B) also indicated that the network of *WAPO1* corresponded to floral transition, in that some gene sets were significantly enriched with reproductive meristem phase change, miR156 miRNA_target_network (miR156 could regulate *TaSPL14* and *TaSPL17* [60, 64]), and SBP TF. After exploring the regulatory regions through motif analysis, we found a novel motif of CArG (CCWWWWWWGG) specifically identified in the promoter region of the *WAPO-A1* gene in AABBDD but not in AABB. We also found that the CArG motif appeared in the genes co-expressed with *WAPO1* (Figure 4).

Previous studies revealed that the promoter of *WAPO1* ortholog contains predicted binding sites for the TFs of MADS-box and SQUAMOSA PROMOTER BINDING PROTEIN (SBP)-like gene [65]. The CArG motif is bound by MADS-box TFs that mainly participated in regulating flowering and floral/spikelet development [66–68]. Flower-specific TFs were confirmed function in removing the H3K27me3 surrounding flower-specific regulatory elements in *Arabidopsis thaliana* [69, 70]. In wheat, the enriched CArG-box motifs were found in the spikelet reduced H3K27me3 peaks [71]. We also found that *WAPO1* is affected by H3K27me3 modification in different developmental stages using data from http://bioinfo.sibs.ac.cn/dynamic_epigenome/. H3K27me3 modification was lower in spikelet I at the booting stage than spikelet II at the flowering and seedling stages, but expression was the opposite. The newly gained CArG motif might have a significant role in the evolution of the *WAPO-A1* gene for functioning in the development of spikelets in hexaploid wheat.

## Discussion and Conclusion

With the latest developments in sequencing technology and assembly methods, many high-quality sequenced genomes of wheat have been produced [72]. The large amounts of data generated require a platform for experimenters to search for genes of interest. Many wheat databases have thus been developed, such as URGI and GrainGenes, which are data repositories of *T. aestivum* and its relatives, and provide tools like BLAST and JBrowse [7, 8]. CerealsDB and Wheat-SnpHub-Portal mainly focus on visualizing SNP data [9, 10]; WheatGmap is committed to analysis of WGS and WES data of *T. aestivum* [11]; and Triticeae-GeneTribe provides homeologous gene relationships among 12 Triticeae species and three out-groups (rice, maize, and *Arabidopsis*) [12]. None of these databases provide a comprehensive search engine for single genes, such as the gene annotation of *WAPO-A1* (Figure 3) provided in ccnWheat. Unlike expression profiling databases such as expVIP, WheatExp, and Wheat eFP Browser (http://bar.utoronto.ca/efp_wheat/cgi-bin/efpWeb.cgi), which only provide expression data of *T. aestivum* [13, 73], ccnWheat integrates RNA-seq data of *T. aestivum, T. dicoccoides, T. urartu*, and *Ae. tauschii* from public platforms, which is useful for studying expression patterns of genes like *WAPO1*. Existing co-expression networks like WheatNet [14], Knetminer [15], and WheatOmics [16] only provide DNA microarray datasets with an early genome assembly version, or just include the co-expression networks of hexaploid wheat.

The ccnWheat database aims to provide an online service platform for comparative analysis of gene function from a multidimensional network across diploid and polyploid wheat species. ccnWheat also includes comprehensive functional annotations (*e.g.*, gene family, GO, miRNA, and metabolic pathways) to predict gene functions. Moreover, ccnWheat includes online tools like GSEA, GO, module, and motif analysis to determine the possible functions of gene sets, and BLAST and ID conversion allow for gene ID conversion between different genome versions. By using ccnWheat, wheat researchers can quickly find key information of a desired gene, predict biological process(es) the gene may participate in, and study the evolutionary history of the gene in wheat of different ploidies.

For genes that have been cloned but whose functional analysis is not very accurate in wheat, ccnWheat can help to further analyze the function. For example, we found that *Arabidopsis UFO* and wheat *WAPO1* have the same 15^th^ amino acid, phenylalanine (F) [55], so we also searched the motif of *UFO*, and identified a variant of the CArG motif in the promoter. Using our UCSC genome browser, we also found that H3K27me3 and H3K4me3 modification have a peak in the gene body of *WAPO1*. However, H3K27me3 modification had obvious differences in different tissues and developmental stages, while H3K4me3 did not. Spikelet reduced H3K27me3 peaks carrying the enriched CArG-box motifs have been found in wheat [71]. Taken together, gaining the CArG motif may affect *WAPO-A1* functions in regulating the total number of spikelets in AABBDD, though this requires further experimental verification. Gene analysis methods like that used for *WAPO1* can also be applied to other cloned genes, to find the evolutionary differences in wheat of different ploidies.

ccnWheat has more possibilities for improvement and we will continue to update it in the future. For example, RNA-seq samples from different growth stages, various stress treatments, and more tissue types can be integrated to build a more robust co-expression network. Concurrently, the 10+ Wheat Genomes project has provided 15 assemblies for different wheat lines from global breeding programs. These wheat accessions and tetraploid durum wheat could be used to analyze co-expression networks and modules with the increase of corresponding RNA-seq samples, so as to more closely link the network with variation and evolution. Epigenomics data like DNase-Seq, ChIP-Seq, ATAC-Seq, MNase-Seq, MeDIP-seq, and BS-seq, which can be used to find peaks with gene expression, can be integrated to clarify the complex relationship between gene expression and chromatin structure. These new additions to ccnWheat will help to mine gene function and breeding in wheat.

By constructing and comparing networks in diploid and tetraploid wheat progenitors, we can dissect the origin and evolution of co-expression networks to better understand the underlying genetic basis for various agronomically important traits of bread wheat. The new ccnWheat platform could facilitate bench scientists identifying key candidate genes for functional studies, and provide genomic scientists a reliable source to decipher key molecular modules during the formation, evolution, and domestication of wheat.

## Data availability

ccnWheat is freely available at http://bioinformatics.cau.edu.cn/ccnWheat.

## CRediT author statement

**Zhongqiu Li:** Data curation, Formal analysis, Visualization, Writing - original draft. **Yiheng Hu:** Data curation, Formal analysis, Writing - review & editing. **Xuelian Ma:** Visualization, Formal analysis, Writing - review & editing. **Lingling Da:** Visualization, Formal analysis. **Jiajie She**: Software, Visualization. **Yue Liu:** Software, Formal analysis. **Xin Yi:** Visualization. **Yaxin Cao:** Software. **Wenying Xu:** Conceptualization, Methodology, Writing - review & editing. **Yuannian Jiao:** Conceptualization, Methodology, Writing - review & editing. **Zhen Su:** Conceptualization, Methodology, Supervision, Project administration, Writing - review & editing. All authors read and approved the final manuscript.

## Acknowledgments

This work was supported by grants from the National Natural Science Foundation of China (Grant Nos. 31970629 and 31771467 for Z. Su, 31870209 for Y. Jiao).

