## Supplemental for "ccnWheat: A Database for Comparing Co-expression Networks Analysis of Allohexaploid Wheat and Its Progenitors"

*Corresponding authors.

**Running title:** *Database for co-expression networks analysis of wheat*

### Case study

#### Using ccnWheat to predict unknown gene function in diploid and polyploid wheat

Cytochrome P450s are the largest family of proteins in higher plants [1], and are known to respond to environmental and developmental cues and tissue and subcellular localization [2]. In the comprehensive comparative analysis of CYP450 in hexaploid wheat and maize, both species have potential functional separation and tissue-specific expression [3]. By comparing the protein sequences of hexaploid AABBDD and its progenitors with wheat protein sequences of CYP450 (http://drnelson.uthsc.edu/cytochromeP450.html), we further identified their P450 domains using PfamScan (https://www.ebi.ac.uk/Tools/pfa/pfamscan/). After filtering, 1564, 823, 462, and 489 full-length *CYP450* genes were found in AABBDD, AABB, AA, and DD, respectively (Table S5). According to the standard naming rules, there are 43, 42, 41, and 43 CYP450 families in AABBDD, AABB, AA, and DD, respectively. In A-type, *CYP71* belongs to the family with the largest number of genes among the four species. In non-A-type, the *CYP96* family accounts for the largest numbers of genes in AABBDD and AA, while the *CYP72* and *CYP51* families account for the largest numbers of genes in AABB and DD, respectively (Figure S7A).

Furthermore, the subgenomes of AABBDD and AABB were classified into A-type and non-A-type families, and the number of A-type gene families was higher than that of non-A-type gene families in all species and corresponding subgenome genes. The gene numbers of A-type and non-A-type families basically conform to the proportion of 1:1:1 or 1:1 in the subgenomes of AABBDD and AABB (Figure S8B). Because both diploids and polyploids contain roots, leaves, and grains, we mainly looked for the expression profile clustering of these three tissue-specific genes. Ultimately, we found 34, five, and three CYP450s in the roots, leaves, and grains, respectively, which were specifically expressed **(**Figure S7-S8**)**. Of these, *CYP71X5* was expressed specifically in grain in all species (Figure S8-S9). *CYP71X5* had five, three, one, and one members in AABBDD, AABB, AA, and DD, respectively (Table S5). There is no corresponding research in *Arabidopsis*. We speculate that *CYP71X5* may be related to grain development.

By searching using the TraesCS6A02G180100 gene (one of the genes of *CYP71X5* in AABBDD) in our database, four of the five genes of *CYP71X5* in AABBDD were included in CFinderM000360 (Figure S9A), which consists of 51 nodes, and the most significant annotation of the module was ‘embryo sac egg cell differentiation’ (GO:0009560) (Figure S9B). Some genes are related to grain development in CFinderM000360: *MOTHER OF FT AND TFL1* (*TaMFT*) has been confirmed to regulate seed germination in wheat [4]; *Prolamin Binding Factor* (*PBF*) may influence embryo size and endosperm starch synthesis [5], *ADP-glucose pyrophosphorylase* (*Agp2*) controls the rate-limiting step in the starch biosynthetic pathway, have Starch accumulation and expression peak at 21 and 15 days post anthesis (DPA) in wheat grains [6, 7], *homogentisic acid geranylgeranyl transferase* (*HGGT*), which catalyzes the committed step of tocotrienol biosynthesis as the primary form of vitamin E in seeds of most monocot plants [8] and *TaKO-B* is related to gibberellin synthesis [9] (Figure S9A). We performed a cis-element analysis of the 3 kb upstream region of all genes in the module, and found that some seed-specific motif such as GCN4_motif and Skn_1_like_motif are significant (Figure S9C**)**. The expression profile of the module shows that the positive genes co-expressed with *CYP71X5* are specifically expressed in grains (Figure S9D**)**. Taken together, *CYP71X5* may be related to grain development, and genes in the CFinderM000360 module may have specific functions in grain development.

### Supplementary material

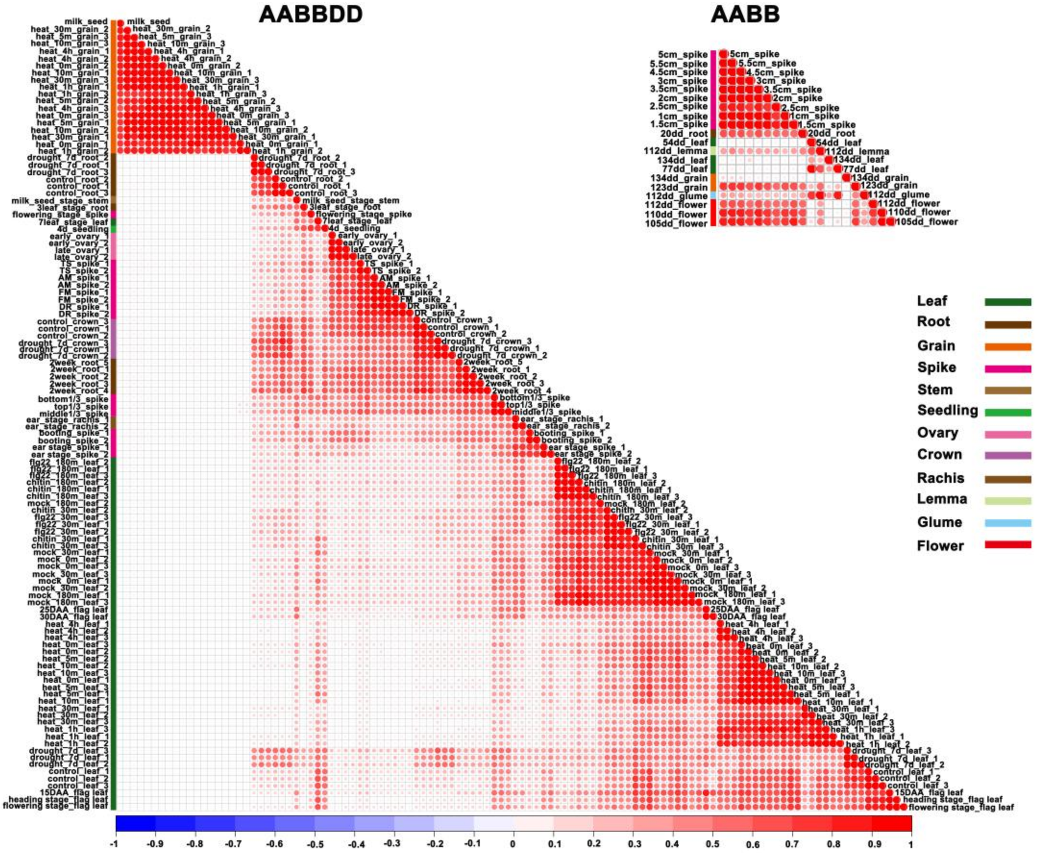


**Figure S1. Correlation analysis for all RNA-seq datasets of AABBDD and AABB**

The diagram shows the correlation and repeatability of samples. The redder the color, the stronger the correlation between the two samples. The bluer the color, the weaker the correlation between the two samples. DR, double-ridge stage; FM, floret meristems; AM, anther primordia stage; TS, tetrads stage; DAA, days after anthesis.

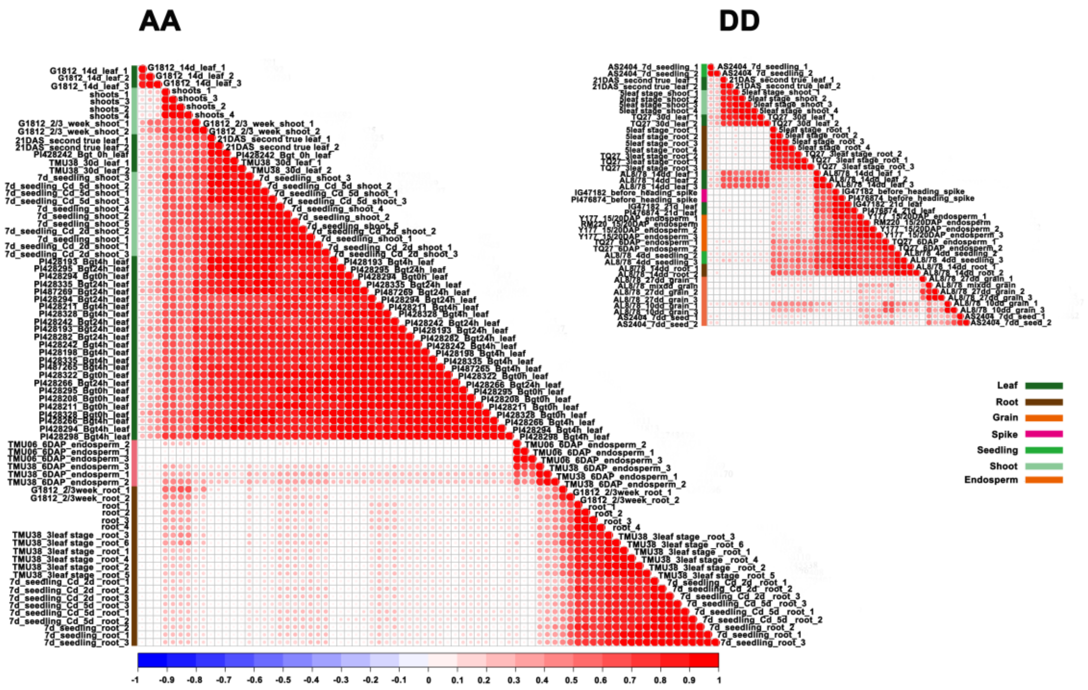


**Figure S2. Correlation analysis diagram for all RNA-seq datasets of AA and DD**

The diagram shows the correlation and repeatability of samples. The redder the color, the stronger the correlation between the two samples. The bluer the color, the weaker the correlation between the two samples. DAS, days after sowing; DAP, days after pollination.

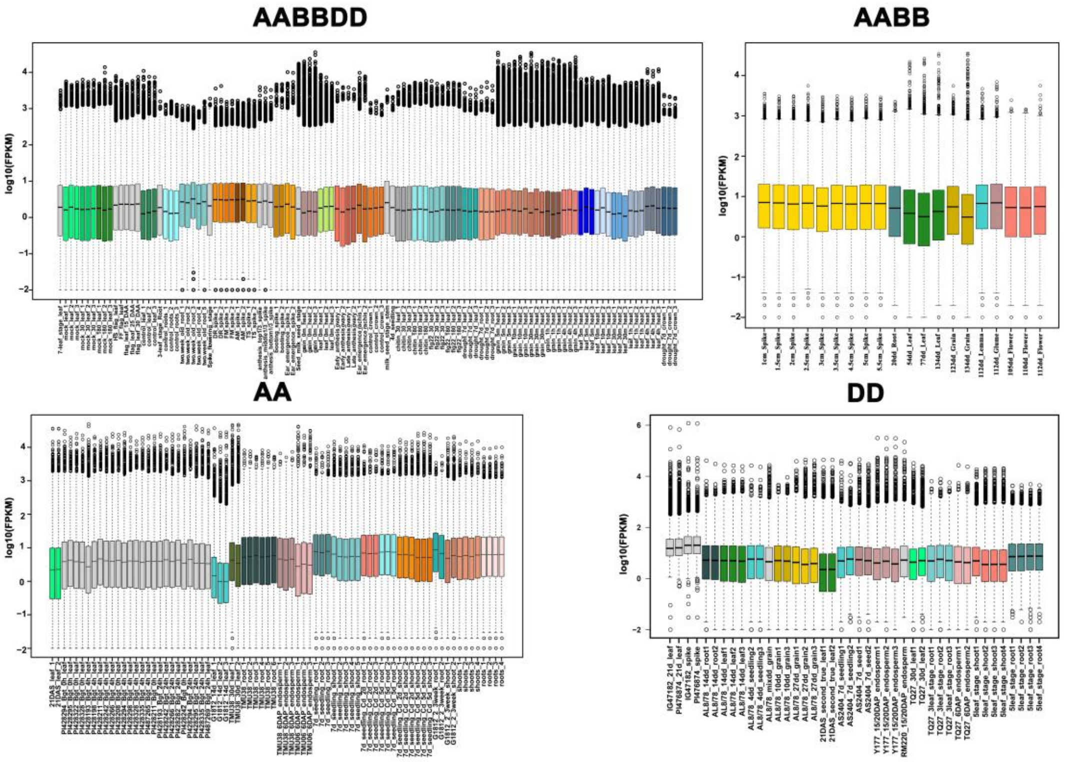


**Figure S3. Boxplots of gene expression values for all species**

The boxplots display the distribution of all genes' FPKM values among the different samples in one species. The x-axis lists the RNA-seq samples, and the y-axis is the gene expression value ($LOG_{10}FPKM$). The same color represents the same tissue or the same duplicate sample. DR, double-ridge stage; FM, floret meristems; AM, anther primordia stage; TS, tetrads stage; DAA, days after anthesis; DAS, days after sowing; DAP, days after pollination.

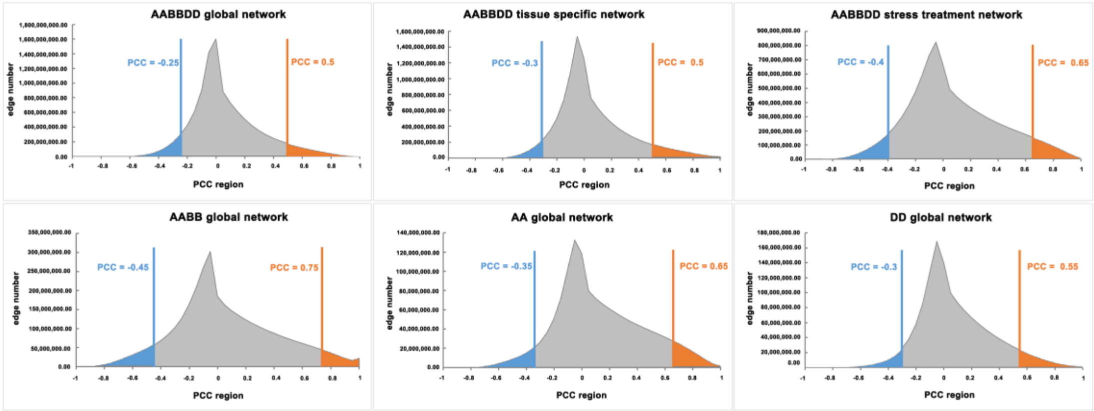


**Figure S4. PCC distributions of all co-expressed gene pairs**

The figure represents the distribution of co-expression relationships in the whole genome. The x-axis represents the region with Pearson correlation coefficient (PCC) from –1 to 1, with an interval of 0.1. The y-axis is PCC value consistent with the interval of co-expressed gene pair (edge). Blue and orange values respectively represent the lowest 5% and highest 5% PCC values, which were set as thresholds, and gene pairs with PCC values in the relevant region were regarded as co-expressed.

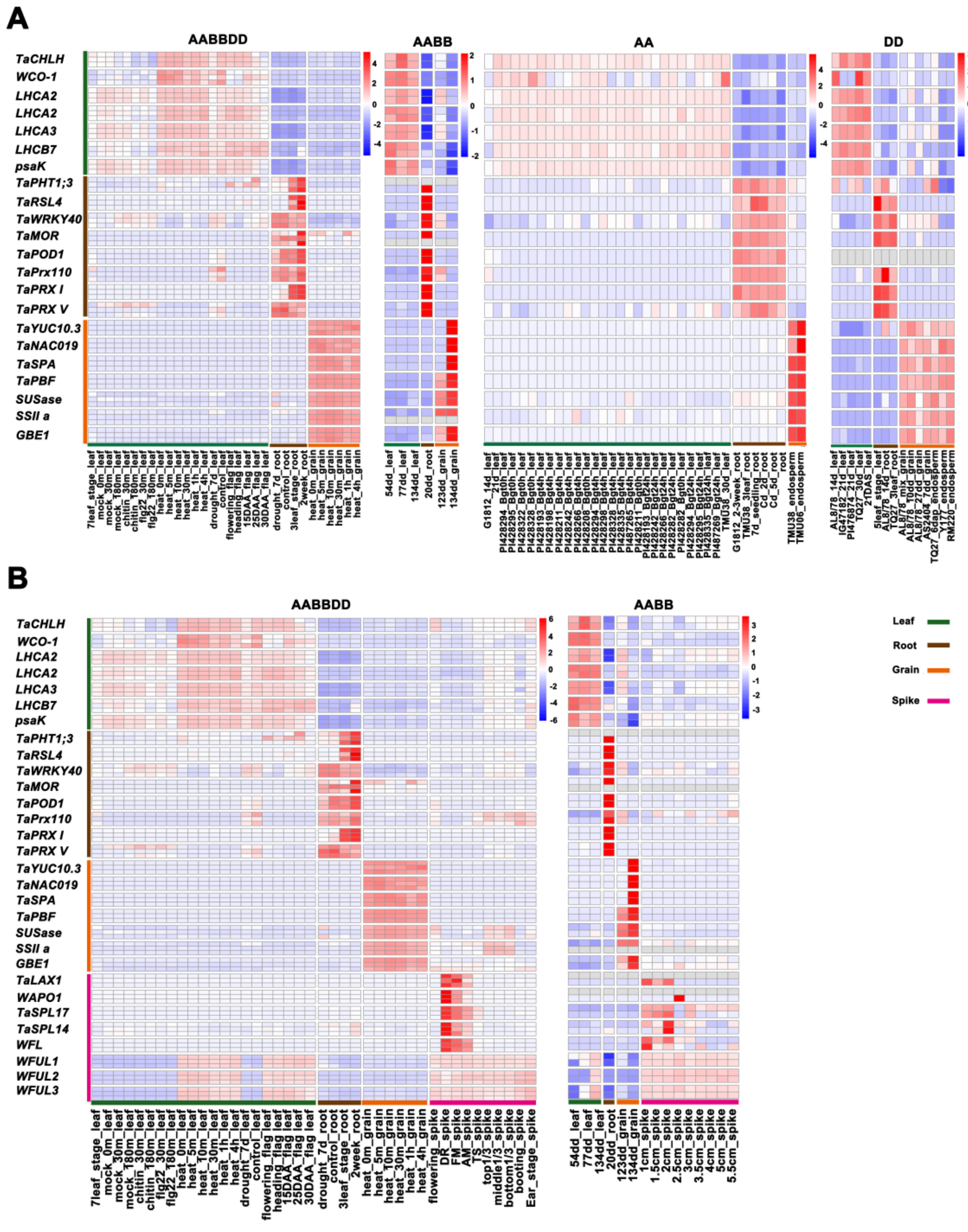


**Figure S5. Analysis of the consistency of expression profiles between species**

**A.** The heatmap shows that genes are specifically expressed in three tissues (leaf, root, and grain) and are basically consistent with their known functions, and their expression trends are consistent in the four species. **B.** The heatmap shows that genes are specifically expressed in four tissues (leaf, root, grain, and spike) and are basically consistent with their known functions, and their expression trends are consistent in the two species. The gene used in the figure collected from the literature or KEGG pathway (gene detail information in Table S4). DR, double-ridge stage; FM, floret meristems; AM, anther primordia stage; TS, tetrads stage; DAA, days after anthesis.

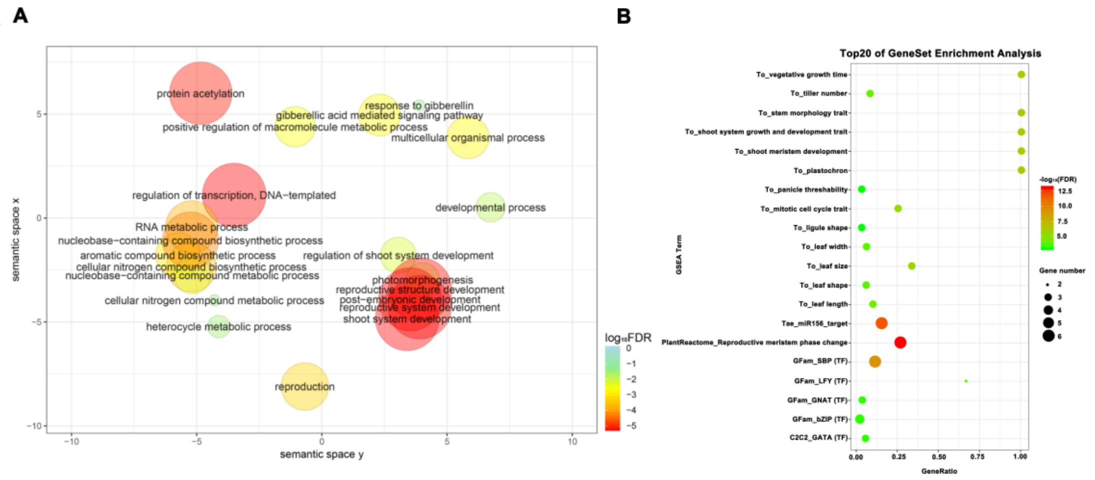


**Figure S6. GO and GSEA analysis of *WAPO1* co-expression network using our database.**

**A.** Gene ontology enrichment analysis of all genes in the *T. aestivum* (AABBDD) global co-expression network for *WAPO1*. **B.** Top 20 most significant gene set enrichment analysis of all genes in the *T. aestivum* (AABBDD) global co-expression network for *WAPO1*.

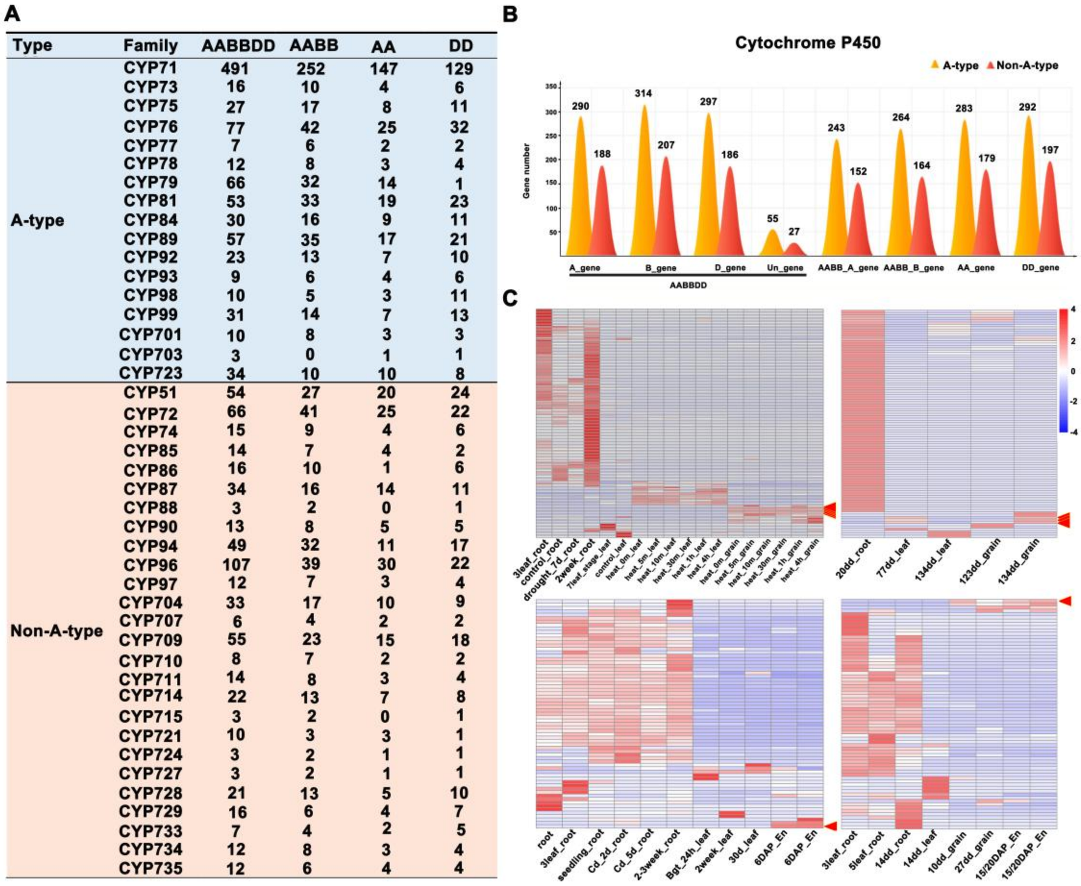

| Type | Family | AABBDD | AABB | AA | DD |
| --- | --- | --- | --- | --- | --- |
| A-type | CYP71 | 491 | 252 | 147 | 129 |
| | CYP73 | 16 | 10 | 4 | 6 |
| | CYP75 | 27 | 17 | 8 | 11 |
| | CYP76 | 77 | 42 | 25 | 32 |
| | CYP77 | 7 | 6 | 2 | 2 |
| | CYP78 | 12 | 8 | 3 | 4 |
| | CYP79 | 66 | 32 | 14 | 1 |
| | CYP81 | 53 | 33 | 19 | 23 |
| | CYP84 | 30 | 16 | 9 | 11 |
| | CYP89 | 57 | 35 | 17 | 21 |
| | CYP92 | 23 | 13 | 7 | 10 |
| | CYP93 | 9 | 6 | 4 | 6 |
| | CYP98 | 10 | 5 | 3 | 11 |
| | CYP99 | 31 | 14 | 7 | 13 |
| | CYP701 | 10 | 8 | 3 | 3 |
| | CYP703 | 3 | 0 | 1 | 1 |
| | CYP723 | 34 | 10 | 10 | 8 |
| Non-A-type | CYP51 | 54 | 27 | 20 | 24 |
| | CYP72 | 66 | 41 | 25 | 22 |
| | CYP74 | 15 | 9 | 4 | 6 |
| | CYP85 | 14 | 7 | 4 | 2 |
| | CYP86 | 16 | 10 | 1 | 6 |
| | CYP87 | 34 | 16 | 14 | 11 |
| | CYP88 | 3 | 2 | 0 | 1 |
| | CYP90 | 13 | 8 | 5 | 5 |
| | CYP94 | 49 | 32 | 11 | 17 |
| | CYP96 | 107 | 39 | 30 | 22 |
| | CYP97 | 12 | 7 | 3 | 4 |
| | CYP704 | 33 | 17 | 10 | 9 |
| | CYP707 | 6 | 4 | 2 | 2 |
| | CYP709 | 55 | 23 | 15 | 18 |
| | CYP710 | 8 | 7 | 2 | 2 |
| | CYP711 | 14 | 8 | 3 | 4 |
| | CYP714 | 22 | 13 | 7 | 8 |
| | CYP715 | 3 | 2 | 0 | 1 |
| | CYP721 | 10 | 3 | 3 | 1 |
| | CYP724 | 3 | 2 | 1 | 1 |
| | CYP727 | 3 | 2 | 1 | 1 |
| | CYP728 | 21 | 13 | 5 | 10 |
| | CYP729 | 16 | 6 | 4 | 7 |
| | CYP733 | 7 | 4 | 2 | 5 |
| | CYP734 | 12 | 8 | 3 | 4 |
| | CYP735 | 12 | 6 | 4 | 4 |



**Figure S7. Cytochrome P450 Classification and specifically expressed P450s.**

**A.** The number of Cytochrome P450 genes in AA, DD, AABB, and AABBDD. Individual subfamilies are missing in a single species, AA lacks two families: CYP88 and CYP715. **B.** Subgenomic genes are divided into A-type and non-A-type. A/B/D_gene means AABBDD subgenome A or B or D gene; AABB_A/B_gene means AABB subgenome A or B gene. **C.** The gene expression profile heatmap displays the specifically expressed P450 genes in roots, leaves, and grain of diploid and polyploid wheat. The red triangle represents the *CYP71X5* gene. En: Endosperm. Detailed information of *CYP71X5* is in Table S5.

**A**

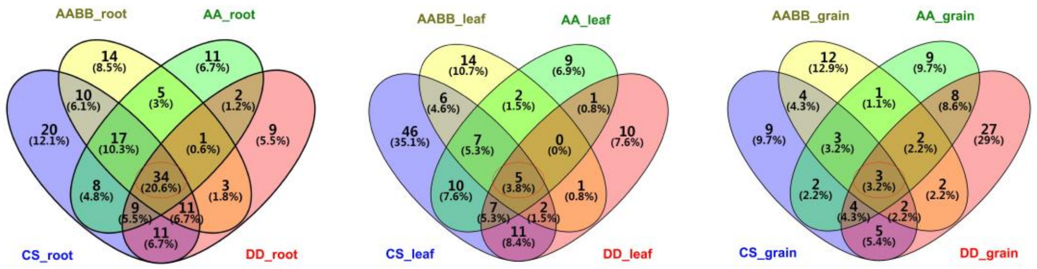


**B**

| Tissue | Common CYP450 |
|---|---|
| root 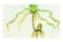 | CYP51H13 CYP51H16 CYP704A41 CYP709C2 CYP709C3v2 CYP710A8 CYP71C101 CYP71E12 CYP71F1 CYP71U4v2 CYP71Y11 CYP723A1 CYP728B3 CYP729A18 CYP72A161 CYP72A167 CYP73A34 CYP75A11 CYP76H21 CYP76H3 CYP76K1 CYP76L1 CYP76L2P CYP81A33 CYP84A5 CYP84A55 CYP87A6 CYP87B11P CYP87C7 CYP89J1 CYP89J2 CYP92A71 CYP94D33 CYP99A31 |
| leaf 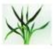 | CYP51G3 CYP96B22 CYP76H21 CYP76M2 CYP71X18 |
| grain 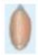 | CYP71X5 CYP71Y12 CYP89YD1 |

**Figure S8. Recognition of CYP450 specifically expressed in roots, leaves, and grains**

**A.** Venn diagram showing the common CYP450 genes specifically expressed in roots, leaves, and grain of diploid and polyploid wheat. **B.** Tissue-specific CYP450 members in all species: 34 members in root, five members in leaf, and three members in grain.

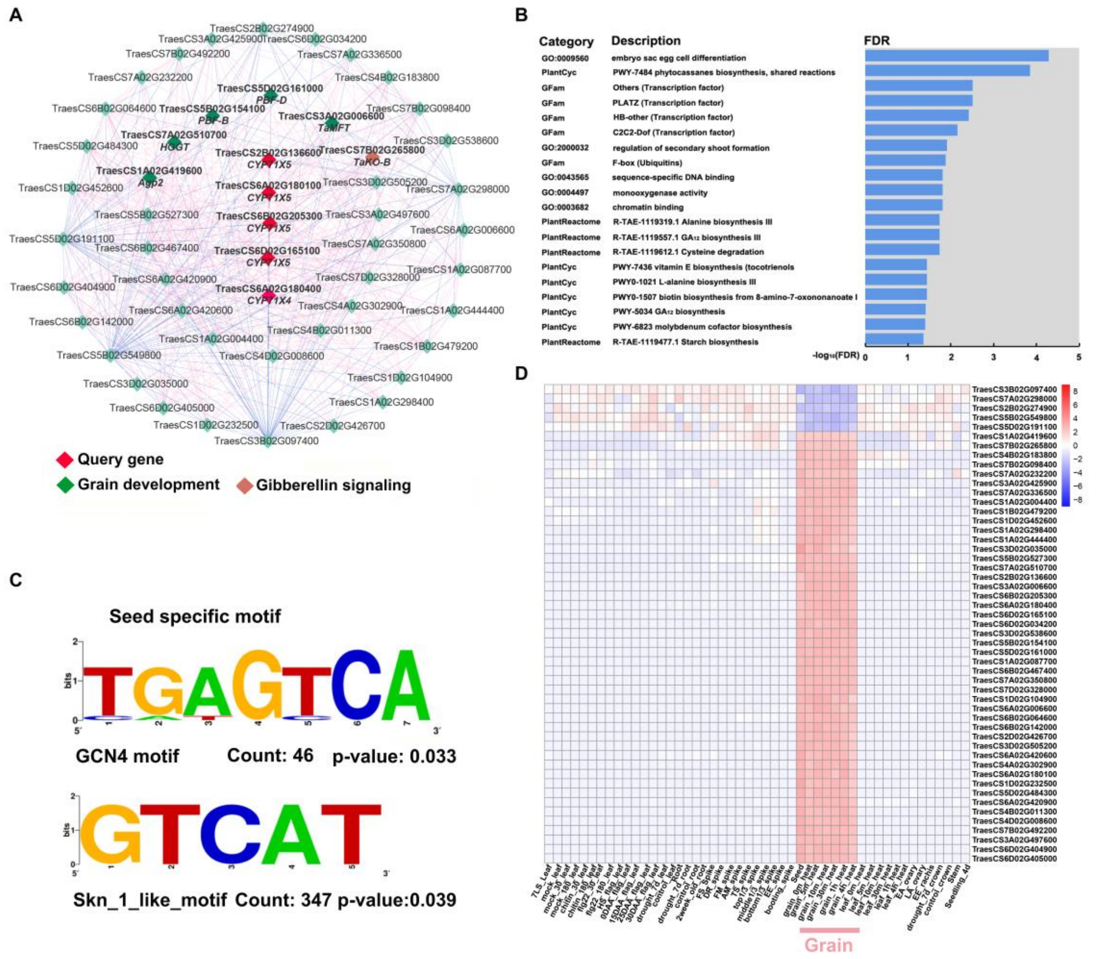


**Figure S9. Analysis of *CYP71X5* module**

**A.** Co-expression module CFinderM000360 identified in T. aestivum (AABBDD). Diamonds represent genes in the module. Red represents *CYP71X5* and *CYP71X4*; green nodes represent genes related to grain development; tangerine nodes represent genes related to Gibberellin signaling. **B.** Module enrichment analysis of CFinderM000360. GO:0009560 embryo sac egg cell differentiation is significantly enriched. Some transcription factors such as PLATZ and HB-other are also significantly enriched. **C.** Cis-elements analysis of CFinderM000360. The seed-specific motif were computed by our custom motif analysis. Count means the number of occurrences of this motif in these genes. **D.** The gene expression profile heatmap displays the gene in the module CFinderM000360. *CYP71X5* module gene is specifically expressed in grain. DR, double-ridge stage; FM, floret meristems; AM, anther primordia stage; TS, tetrads stage; DAA, days after anthesis; LS, leaf stage; HS, heading stage; FS, flowering stage; EE, ear emergence; EA, early anthesis; LA, late anthesis.

**Table S1. Details of RNA-seq sample resources**

**Table S2. Data information and mapping ratios of RNA-seq**

**Table S3. Data information and mapping ratios of epigenetics in AABBDD**

**Table S4. Information of gene collected from the literature and KEGG pathway**
